# Age-related macular degeneration affects the optic radiation white matter projecting to locations of retinal damage

**DOI:** 10.1101/336206

**Authors:** Shoyo Yoshimine, Shumpei Ogawa, Hiroshi Horiguchi, Masahiko Terao, Atsushi Miyazaki, Kenji Matsumoto, Hiroshi Tsuneoka, Tadashi Nakano, Yoichiro Masuda, Franco Pestilli

## Abstract

**Purpose:** We investigated the impact of age-related macular degeneration (AMD) on visual acuity and the visual white matter.

**Methods:** We combined an adaptive cortical atlas and diffusion-weighted magnetic resonance imaging (dMRI) and tractography to separate optic radiation (OR) projections to different retinal eccentricities in human primary visual cortex. We exploited the known anatomical organization of the OR and clinically relevant data to segment the OR into three primary components projecting to fovea, mid- and far-periphery. We measured white matter tissue properties – (fractional anisotropy, linearity, planarity, sphericity) along the aforementioned three components of the optic radiation to compare AMD patients and controls.

**Results:** We found differences in white matter properties specific to OR white matter fascicles projecting to primary visual cortex locations corresponding to the location of retinal damage (fovea). Additionally, we show that the magnitude of white matter properties in AMD patients’ correlates with visual acuity. In sum, we demonstrate a specific relation between visual loss, anatomical location of retinal damage and white matter damage in AMD patients. Importantly, we demonstrate that these changes are so profound that can be detected using magnetic resonance imaging data with clinical resolution. The conserved mapping between retinal and white matter damage suggests that retinal neurodegeneration might be a primary cause of white matter degeneration in AMD patients.

**Conclusions:** The results highlight the impact of eye disease on brain tissue, a process that may become an important target to monitor during the course of treatment.

## Introduction

Human aging can be associated with degeneration of both or either the retinal pigment epithelium and photoreceptor cells. Such degeneration often starts at the center of the retina in the *Macula*, a clinical syndrome referred to as age-related macular degeneration (AMD). AMD often results in irreversible loss of visual function. By affecting more than 25% of individuals older than 75 years (Ratnapriya and Chew 2013), AMD has been prospected to become a disease of global impact with a total number of individuals affected by 2020 reaching 196 millions worldwide (Wong et al. 2014).

Modern technologies for mapping the human brain allow investigators to measure both micro- and macroscopic properties of the white matter tissue organization and composition *in vivo,* these methods have the potential to diagnose and track disease progression across the lifespan. Diffusion-weighted magnetic resonance imaging (dMRI) and computational tractography are two methods that, when combined, can map the macroscopic anatomical organization as well as the microscopic tissue properties of the human white matter (Jbabdi et al. 2015; Wandell 2016; A. Rokem et al. 2017).

Measurements using dMRI have shown biological changes in white matter associated with neurodegeneration. For example, the human white matter is affected during normal aging as well as in age-related diseases, such as Alzheimer’s or progressive cognitive degeneration (Mayda et al. 2009; Wang et al. 2010, 2012; Yeatman, Wandell, and Mezer 2014; Thomason and Thompson 2011). The visual white matter pathways have been also shown to be affected by visual and eye disease, for example, developmental prosopagnosia (Gomez et al. 2015a; Thomas et al. 2008), amblyopia (Allen et al. 2015a, 2018; Duan et al. 2015a), and retinal ganglion-cell damage (Ogawa et al. 2014) have all been reported to affect properties of the human white matter, please see (Ariel Rokem et al. 2016; Millington, Ajina, and Bridge 2014) for reviews. Evidence from *in vivo* studies suggest that macular degeneration also affects volume of the human primary visual cortex and the visual pathways (Prins et al. 2016; Prins, Hanekamp, and Cornelissen 2016; Hernowo et al. 2014a; Malania et al. 2017). Yet to date, the complete set of mechanisms by which AMD has long term effects on the human brain and behavior are not well understood. Importantly, the degree to which the changes in visual white matter can inform AMD diagnose – i.e., classify individuals as AMD patient or within the normal aging population – has not been established. We address both this points.

We estimated the properties of the visual white matter using multiple models of tissue microstructure – fractional anisotropy (Basser and Pierpaoli 1998a), planarity, and sphericity (Carl-Fredrik Westin et al. 1997; C-F Westin et al. 2002). Critically, we show changes in white matter tissue properties within the optic radiation that are specific to fascicles projecting from central, foveal locations, those with retinal damage. To do so, we performed an advanced, anatomically precise characterization of the relationship between the locus of retinal damage in AMD (fovea) and the white matter fascicles within the optic radiation projecting to locations of primary visual cortex that represent different eccentricities – fovea, parafoveal and peripheral regions combining anatomical and advanced computational methods (Benson et al. 2012, 2014). This fine grade mapping between the retinotopic location of damage and white matter goes beyond previous reports (Prins et al. 2016; Prins, Hanekamp, and Cornelissen 2016; Hernowo et al. 2014b; Malania et al. 2017). Furthermore, we demonstrate that changes in the microstructural properties of the OR correlate with AMD patients’ visual acuity, indicating a relationship between the tissue properties of the OR and the long-term loss in visual function that characterizes AMD. These results show a conserved mapping between the location of retinal damage and white matter fascicles, suggesting that retinal neurodegeneration might cause long-term changes in the visual white matter of the OR in AMD patients.

## Methods

This research was approved by the ethical committees of the Jikei University School of Medicine, Tamagawa University, and Atsugi City Hospital. All participants provided written informed consent to participate in the project. All procedures conformed to the Declaration of Helsinki.

Two experienced ophthalmologists (one was the first author) diagnosed AMD at the Jikei University School of Medicine or at the Atsugi City Hospital. All subjects were given an ophthalmological examination, including best-corrected visual acuity, intraocular pressure, slit lamp microscopy, fundus examination, and optical coherence tomography (OCT) measurements. To do this, we first measured visual acuity, which is a perception of minimum angle of resolution, in patients and control subjects using a standard Landolt-C stimulus. Subjects performed a 4-alternative-forced-choice task by indicating the side toward which the “C” of the Landolt stimulus faced (top, bottom, left or right). For statistical analysis, visual acuity was converted to a logarithmic minimum angle of resolution (logMAR).

Eight patients suffering from AMD (seven males; mean age 74.8 years, range 62-84 years). Six of them had bilateral AMD. The others had unilateral damage. All AMD subjects were undergoing anti-vascular endothelial growth factor therapy (anti-VEGF; (D. M. Brown et al. 2006; Heier et al. 2012; Krüger Falk et al. 2013). Twelve control subjects (six males; mean age 65.3 years, range 59-74 years) with normal, or corrected to normal, visual acuity. None of the patients or controls had a history of brain disease. See Table 1 for additional details about the patients’ and controls’ biographical and biometric information.

**Table 1.**
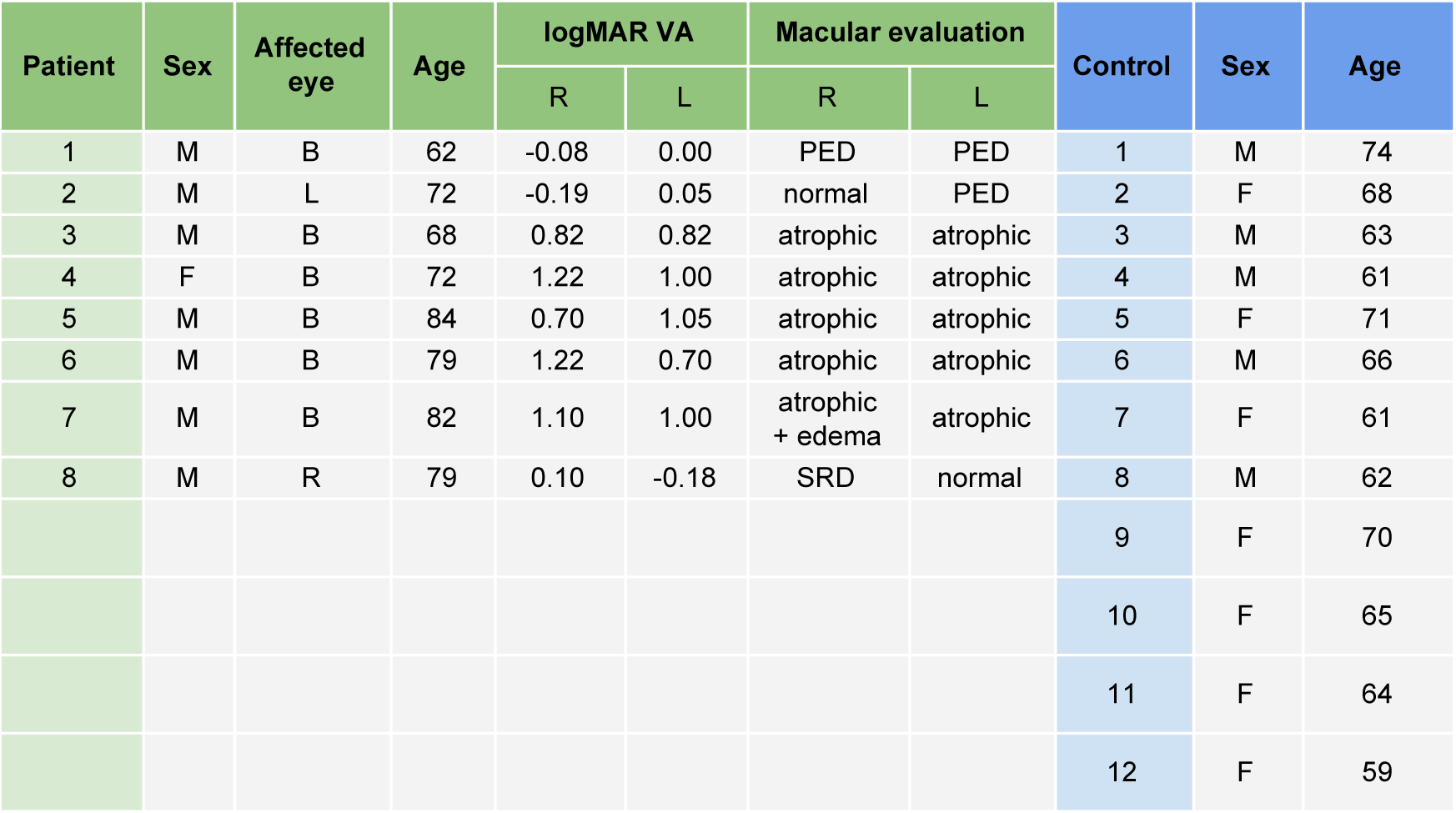
Biographic and biometric measurements of the AMD patients and control participants. The *logMAR VA* measurements indicate the minimum angle of resolution visual acuity in log units. PED stands for pigment epithelial detachment, SRD stands for serous retinal detachment. Green columns report information about patients; blue, control subjects. Letters B, R and L stand for bilateral, right and left eye, respectively.

| Patient | Sex | Affected eye | Age | logMAR VA |  | Macular evaluation |  | Control | Sex | Age |
| --- | --- | --- | --- | --- | --- | --- | --- | --- | --- | --- |
|  |  |  |  | R | L | R | L |  |  |  |
| 1 | M | B | 62 | -0.08 | 0.00 | PED | PED | 1 | M | 74 |
| 2 | M | L | 72 | -0.19 | 0.05 | normal | PED | 2 | F | 68 |
| 3 | M | B | 68 | 0.82 | 0.82 | atrophic | atrophic | 3 | M | 63 |
| 4 | F | B | 72 | 1.22 | 1.00 | atrophic | atrophic | 4 | M | 61 |
| 5 | M | B | 84 | 0.70 | 1.05 | atrophic | atrophic | 5 | F | 71 |
| 6 | M | B | 79 | 1.22 | 0.70 | atrophic | atrophic | 6 | M | 66 |
| 7 | M | B | 82 | 1.10 | 1.00 | atrophic + edema | atrophic | 7 | F | 61 |
| 8 | M | R | 79 | 0.10 | -0.18 | SRD | normal | 8 | M | 62 |
|  |  |  |  |  |  |  |  | 9 | F | 70 |
|  |  |  |  |  |  |  |  | 10 | F | 65 |
|  |  |  |  |  |  |  |  | 11 | F | 64 |
|  |  |  |  |  |  |  |  | 12 | F | 59 |

Structural evaluation of the retina was performed using optical coherence tomography (Cirrus HD-OCT; Carl Zeiss Meditec, Dublin, CA, USA). A typical fundus and optical coherence tomography (OCT) image in patient with AMD is shown in **Figure 1**. The OCT data distinguish between patients with other different diseases.

**Figure 1.**
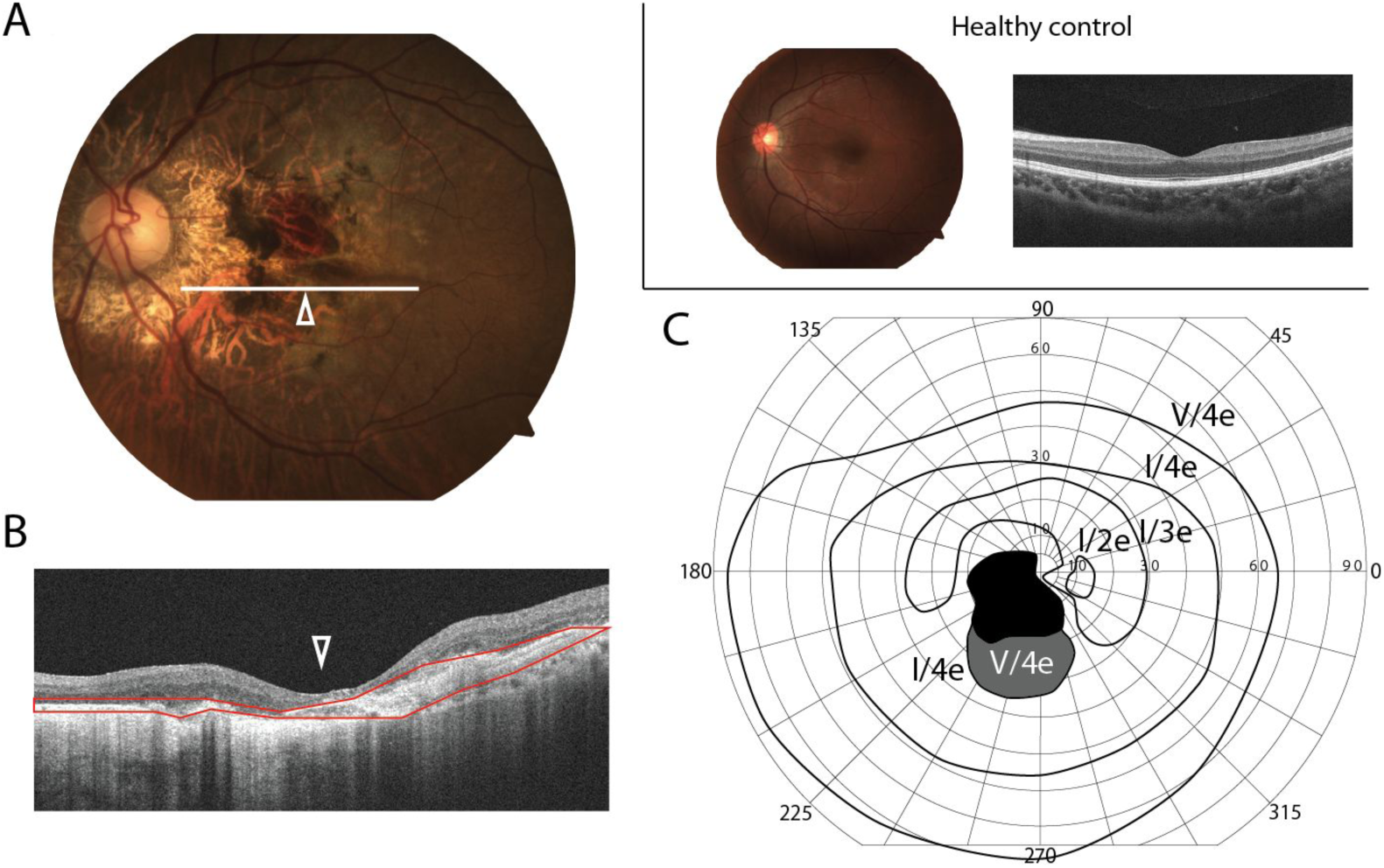
Measurements of retinal damage. Measurements are reported for a single left-eye in **A, B**, and **C. A**. Example fundus (retinal image). Left-hand panel. The white horizontal line indicates the approximate location of the optical coherence tomography measurement (OCT; **Figure 1B**). The white triangle marks the center of the fovea. Right-hand panel. Fundus and OCT image for one example control subject. **B**. OCT image. Section of the retina through the white line in **A**. The red region indicates the retinal area with AMD-related tissue degeneration. The structure of outer retina layer is almost completely degenerated. The inner / outer segment line is invisible, indicating photoreceptor damage. White triangle indicates the fovea as in **A**. **C**. Visual field, as measured by Goldmann perimetry. Black depicts regions of absolute scotoma. The participant could not detect even the largest and brightest stimuli in Goldmann perimetry set. Paracentral gray indicates retinal areas where the patient could detect only the largest and brightest stimulus (V/4e) in the Goldmann perimetry set. Other regions mark locations with either limited damage or no damage.

## Visual field measurement

Visual field measurements were acquired in six patients using standard Goldmann perimetry. Two patients did not consent to receive the perimetry test. The distribution of visual sensitivity across the visual field differed across subjects. We used kinetic targets to define the absolute scotoma as the region in which subjects could not detect the highest-contrast and largest-size stimuli V/4e; 64 mm^2^ (visual angle 1.72° diameter), 318 cd/m^2^. Relative scotoma was defined as the region in which subject could not detect the stimuli I/2e and I/3e; 0.25 mm^2^ (visual angle 0.11°). Five eyes had relative scotoma and five eyes had absolute scotoma. Two eyes were normal (see **Figure 1c** for example measurements). The position of the (absolute or relative) scotoma was recorded in all patients closest to macula (center of the visual field).

### Magnetic resonance image data acquisition

Magnetic resonance imaging (MRI) data acquisition and preprocessing methods used in this study were identical to those developed for (Ogawa et al. 2014). Below we briefly describe the major details of MRI measurements and analyses.

All data were acquired at the Brain Science Institute of Tamagawa University, Tokyo, Japan. High-resolution T1-weighted anatomical images were acquired for each subject using a three-dimensional spoiled-gradient recalled acquisition (SPGR; 1×1×1 *mm* voxel size; scan duration 9 minutes, 18 seconds) with sagittal plane acquisition on 3T Siemens MAGNETOM Trio Tim scanner (Siemens, Erlangen, Germany) using a 12-channel head coil.

Diffusion-weighted magnetic resonance images were acquired using two repetitions of a single-shot spin-echo planar imaging sequence (93 ms echo time (TE); 7.5 seconds repetition time (TR); 230 *mm* field of view; 230 × 56 matrix size; ±1562 Hz/Px bandwidth; 317s duration). We acquired 56 axial, 1.8-mm-thick slices (no gap) for two b-values, b = 0 and b = 1000 s/mm^2^. The b=1000 data were obtained by applying gradients along 12 unique diffusion directions.

### Data preprocessing

Anatomical images were aligned to the anterior commissure–posterior commissure (AC-PC) plane using a 6° rigid-body transformation. Diffusion-weighted images were corrected for eddy-currents, motion compensated, and aligned to the high-resolution anatomical images using a 14-parameter constrained nonlinear coregistration. All diffusion-weighted images were resampled to 2 *mm* isotropic resolution using trilinear interpolation (Ashburner and John 2009, 2012). An eddy-current intensity correction was applied to the diffusion-weighted images at the resampling stage. Diffusion-weighting gradient directions were reoriented by applying the same transformation used on the diffusion-weighted images.

#### Diffusion tensor model

Fractional anisotropy, mean, radial and axial diffusivity were estimated by fitting a diffusion tensor model to each voxel signal (Basser and Pierpaoli 1998a). Fractional anisotropy (FA) was defined as the normalized standard deviation of the three eigenvalues of the diffusion tensor and indicates the degree to which the diffusion ellipsoid is anisotropic (i.e., one or two eigenvalues are larger than the mean of all three eigenvalues). Mean diffusivity (MD) was defined as the mean of the three eigenvalues. Axial diffusivity (AD) was defined as the apparent diffusion coefficient measured along the principal axis of the tensor model in each voxel. Radial diffusivity (RD) was defined as the average of the diffusivity in the two minor axes of the tensor model. All image analysis software is open source and published at https://github.com/vistalab/vistasoft (Ariel Rokem et al. 2015; Pestilli, Franco, et al. 2014; Dougherty et al. 2007).

#### Brain segmentation and regions of interest definition

The brain cortical-surface and optic chiasm were reconstructed and segmented using FreeSurfer(Bruce Fischl 2012); version 5.1.0; (http://surfer.nmr.mgh.harvard.edu). We used a probabilistic anatomical atlas to estimate the location of primary visual cortex (Bruce Fischl 2012; B. Fischl et al. 2007; Hinds et al. 2008). The approximate location of the lateral geniculate nucleus (LGN) was estimated manually on the high-resolution anatomical image (Ogawa et al. 2014b; Allen et al. 2015b; Millington, Ajina, and Bridge 2014).

### Anatomy of white matter tracts and relation to ophthalmic examination

We identified the optic tract (OT) and radiation (OR) using probabilistic fiber tractography which has been shown to provide improved results for mapping white matter structures (Pestilli, Franco, et al. 2014; Sherbondy et al. 2008). All methods were reported in previous publications (Sherbondy et al. 2008; Ogawa et al. 2014).

Diffusion properties (Fractional Anisotropy, FA; Mean Diffusivity, MD; Radial Diffusivity, RD; Axial Diffusivity, AD) were calculated separately at 100 locations along the length of the OT and OR, these measurements hereafter are referred to as tract profiles (Yeatman et al. 2012; Rykhlevskaia et al. 2009). This visualization enables us to compare the diffusion properties of individual subjects with controls along the full path of the OT and OR.

We compared the OR tract profiles in individual AMD patients with the distribution of tract profiles in the control groups. To do so, we computed mean and standard deviation of the diffusion properties at each one of the 100 locations in the OT and OR. We excluded the first and last 10% of these locations leaving us with 80 measurements per white matter tract. **Figure 2** shows the measurements. Statistical analyses were performed by computing a non-paired t-test along these tract profiles as described in previous studies (Yeatman et al. 2012; Ogawa et al. 2014; Allen et al. 2015b; Rykhlevskaia et al. 2009; Ajina et al. 2015a; Leong et al. 2016a; Gomez et al. 2015b).

**Figure 2.**
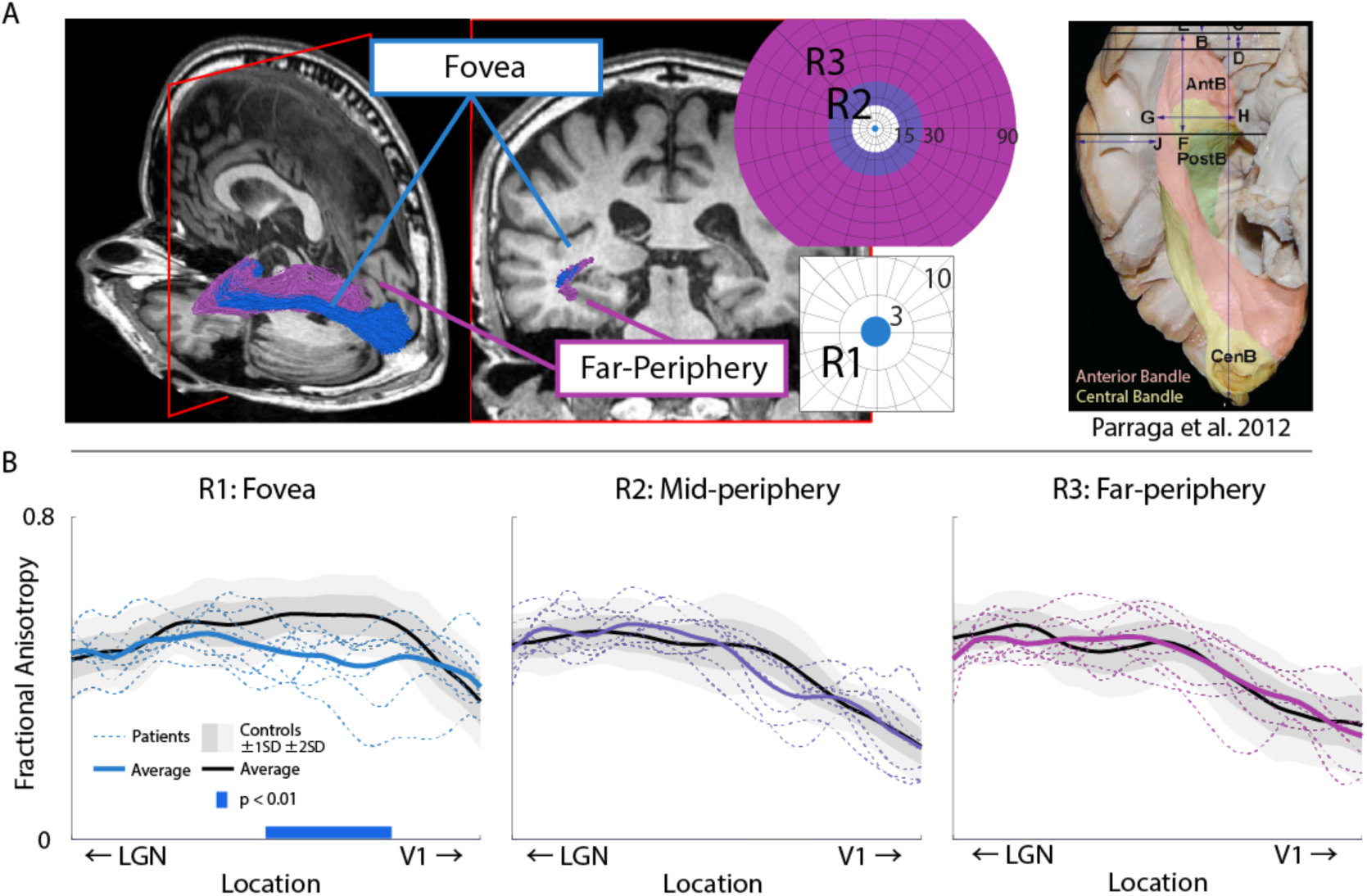
White matter properties along optic radiation grouped by V1 field-map eccentricity. A. We combine primary visual cortex surface topology and fascicles properties to separate white projections to fovea and periphery. *Left-hand panel*. The anatomical organization of the optic radiation (OR) fascicles showing spatially separated fascicles projecting to foveal and peripheral visual cortex. *Right-hand panel.* We show both a *post mortem* anatomical segmentation depicting the known separation of the OR fascicles bundles projecting to fovea and periphery. The two bundles are generally referred to as the central bundle (yellow) and anterior bundle (salmon). Combining V1 surface topology and white matter fascicles length properties separates bundles projecting primarily to fovea (blue, R1) and periphery (purple, R3). **B. Tract profiles of white matter summary statistics**. Fractional anisotropy (FA) is plotted along the length of the optic radiation for AMD patients and control participants for white matter bundles projecting to different visual field locations. Individual patients, dashed lines. Group averages, thick lines, black control group. Shaded areas ±1 and 2 standard deviations (SD) in the control group. Locations of significant difference, p<0.01, between patient and control groups are highlighted by a solid blue bar at the bottom of the plot. We found that only about 21% of the white matter voxels were shared between the *fovea (R1), mid-* and *far-periphery* (R2 and R3 respectively).

#### Visual field maps and white matter

In addition to identifying the full anatomy of the OT and OR, we also identified the relation between the terminations of the OR fascicles and visual field eccentricity in V1. It is established that human V1 is organized into a retinotopic-conserving map structure (Wandell, Dumoulin, and Brewer 2007; Wandell and Smirnakis 2010; Wandell and Winawer 2015, 2011). This means that nearby locations in the visual environment map to nearby locations in the retina and in V1.

It has been shown that the retinotopic organization of V1 can be mapped using anatomical methods(Benson et al. 2012). We used the atlas by Benson and colleagues to identify three regions within V1 of each participant in the study. V1 was subdivided into three regions (R) depending on their putative visual field eccentricity mapping. The three regions mapped from central fovea to far periphery with predicted eccentricity values divided as follows: R1, 0-3°; R2, 15-30°; R3, 30-90°. We used these three V1 regions to subdivide the white matter fascicles of the OR into three groups with fascicles each terminating into different V1 regions R1, R2 and R3. This was possible because both the length and termination of white matter fascicles differed depending on their projection to V1, see **Figure 3A**. Given the resolution of the dMRI data, the spatial uncertainty of tractography to accurately map fascicles within individual voxels and the uncertainty added by the mapping between anatomical atlas to each individual brain, the effective eccentricity mapping between R1, R2 and R3 might be slightly different than the nominal one. For this reason we refer to R1 as *Fovea*, R2 as *Mid-periphery* and R3 as *Far-periphery* regions.

**Figure 3.**
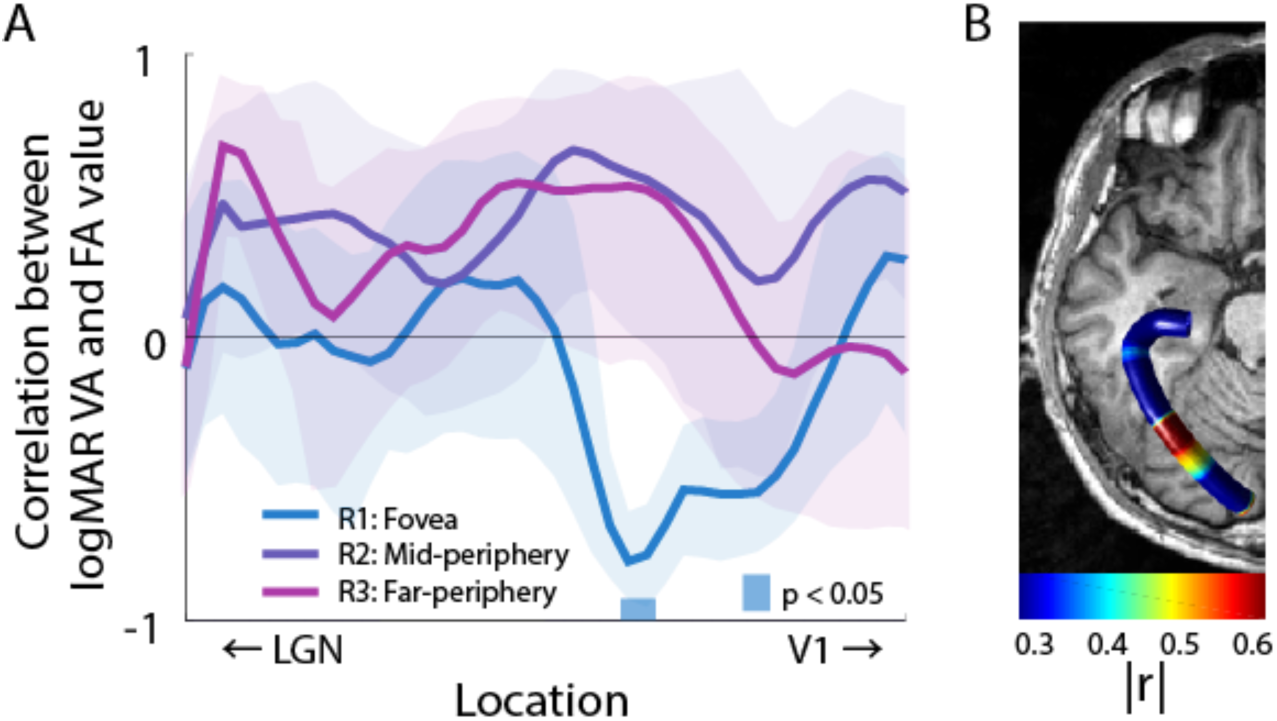
Correlation analysis between visual acuity and white matter damage. **A**. FA values along the OR correlated with participants’ logMAR visual acuity at the location of white matter damage. Shaded areas show 80% confidence interval. Light blue band above the abscissa shows location of statistically significant correlation between logMAR visual acuity and FA. Darker blue band below abscissa reports locations of significantly different FA values between AMD and control subjects (see Figure 2B for reference). **B**. Anatomical position of the results. Average tract profile displaying the correlation between behavioral acuity and FA along the anatomy of the OR. All data displays estimates for the foveal region (R1). Two nodes in the core white matter of the OR were found to be statistically significant with p<0.05.

#### Visual acuity measurements and white matter

We investigated the relation between white matter properties in the OR and behavioral measurement of visual acuity. To do so we used a standard Landolt-C ophthalmologic test to measure visual acuity at either fovea or at the preferred retinal locus of each individual patient – no measurements were taken in control subjects. We correlated the visual acuity measurement with those in white matter properties in each participant. White matter measures (FA, MD, AD and RD) were averaged across both for this analysis, **Figure 3** shows the results.

## Results

### Visual white matter and retinal damage

Goldmann Perimetry measurements revealed absolute scotomas in the most central retinal locations of AMD patients, with improved retinal function at higher eccentricity (**Figure 1**). We measured the anatomical shape and position of the optic tract (OT) and optic radiation (OR) in all patients and control subjects (**Figure 2** and **Supplementary Figure 1-2**). We measured the white matter tissue along the OT and OR using tract profiles (Yeatman et al. 2012; Ogawa et al. 2014; Allen et al. 2015a; Ajina et al. 2015b; Duan et al. 2015a; Gomez et al. 2015a; Leong et al. 2016b; Libero et al. 2016) and the tensor model (Basser and Jones 2002; Basser and Pierpaoli 1998b). **Supplementary Figure 1A** shows mean and individual FA tract profiles of AMD patients (red) as well as the mean across control subjects (black) in the OT. Measurements show no statistically significant difference in FA between AMD patients and controls in the optic tract (**Supplementary Figure 1A**). This is in contrast with a statistically significant difference in FA in the OR (**Supplementary Figure 1B**, orange bar; p<0.01).

Because of the stereotyped location of retinal damage in the fovea of the AMD patients, we predicted that the white matter tissue of optic radiation (OR) would be affected primarily in fascicles projecting to the central visual field. To test this prediction, we subdivided the OR into three branches projecting to Foveal (R1), Mid-periphery (R2) and Far-periphery (R3) locations (**Figure 2**). To do so, we used a recent automatic retinotopic mapping methods (Benson et al. 2014, 2012) and the known anatomical subdivision of the OR (**Figure 2A** and **B**) to separate the white matter projecting to the three eccentricity ranges of primary visual cortex (V1; see **Methods** for additional details).

Fractional anisotropy (FA), mean diffusivity (MD), axial diffusivity (AD) and radial diffusivity (RD) were measured for each participant along the OR. **Figure 2B** show mean and individual FA tract profiles of AMD patients (red), as well as the mean across control participants (black). OR FA measures differed between patients and controls. Mean FA values within the central portion of the OR (core white matter) were significantly different between patients and control subjects (t-test, p < 10^-34^). Furthermore, FA was lower in AMD patients at a specific location along the optic radiation (**Figure 2B**, left-hand panel; p<0.01, t-test). **Supplementary Figure 2A-C** shows the results for MD, AD and RD. We went beyond the tensor model to show that the measured effect is robust to the choice of model and parameters used for characterizing the diffusion signal.

We further tested the specificity of the effect in the OR to demonstrate that the result is not driven by other differences in the groups of controls and patients. To do so, we mapped eleven major human white matter tracts, namely the Inferior and Superior Longitudinal Fasciculi (ILF and SLF, respectively), the Arcuate Fasciculus, the Uncinate Fasciculus, the Inferior Fronto Occipital Fasciculus (IFOF), Cingulum Hippocampus and Cingulate, the Callosum Forceps Major and Minor, Corticospinal Tract and the Thalamic Radiation (Caiafa and Pestilli 2017; Pestilli, Yeatman, et al. 2014; Yeatman et al. 2012). **Supplementary Figure 2D** reports the tract profiles for all these tracts (average between the two hemispheres, except Forceps Major and Minor). We compared the FA values in each of these tracts between control and patients groups. A total of 440 (40 nodes in 11 tracts) were compared, no statistical difference was found in the majority of the white matter in these tracts, with the exception of 15 nodes (p<0.01; five in the Cingulum, five in the IFOF, two in the Callosum Forceps Major and three in the Callosum Forceps Minor). This is small number of nodes passing significance is expected given the large number of comparisons (440). The lack of reliable differences in FA in the major white matter tracts suggests that the strong differences in FA in the OR might be due to the relation between retinal damage and the visual white matter, and not by generalized brain differences between the two groups. To further, investigate this, next we compared whether a relation exists between individual subjects Visual Acuity (VA) and the FA in the OR.

### Visual white matter and behavior: Fractional anisotropy of white matter projecting to fovea predicts visual acuity

We established the association between patients’ logMAR visual acuity (hereafter only referred to as “visual acuity”) and white matter properties of the OR. We predicted that FA values in individual patients would be associated with visual acuity; such that the worse the visual acuity the lower FA. We tested this prediction by measuring best corrected visual acuity in each patient using a standard Landolt-C ophthalmologic test (see **Methods**).

In sum, these results show a fine-scale anatomical mapping between location of putative neurodegeneration, retinal scotoma and behavioral impairment. The result is consistent with previous reports of correlation between FA and visual acuity in other types of visual disease, such as retinitis pigmentosa (Ohno et al. 2015), and suggests that the magnitude of FA along the optic radiation may be an important predictor of visual impairment in AMD patients.

### Visual white matter and disease diagnose: The tissue properties of the optic radiation predict subjects class membership

Diagnosing visual disease from brain tissue properties and investigating the relation between brain and eye disease is critical to advancing a comprehensive understanding of the visual disease (Wandell and Le 2017). Here, we were interested in understanding the degree to which the white matter properties of the optic radiation to predict the subjects class membership. For this reason, we performed two additional analyses. First, we used the measured properties of white matter tissue microstructure (FA) at fifty locations along the optic radiation and performed a logistic regression analysis (Hastie, Tibshirani, and Friedman 2013; Green and Swets 1988) to separate the subjects into two groups, patients or controls (**Figure 4A)**. Logistic regression measures the Area Under the Curve (AUC) to indicate classification performance. Our analysis compared two models; (1) One model assumed that patients and controls groups were different and (2) another in which they were assumed to not be different. Higher AUC values indicated that the model separating subjects groups (model 1) has a higher statistical power than other one (model 2; (Hastie, Tibshirani, and Friedman 2013; Green and Swets 1988). Results demonstrated that FA in specific location along the OR allowed distinguishing between the two patient groups (Figure 3B). Furthermore, FA was more predictive of subjects group for fascicles projecting to foveal V1 (max AUC R1: 0.979, R2: 0.802, R3: 0.791). To establish whether all segments of the OR (R1, R2 or R3) predicted class membership or just the segment projecting to the location of retinal damage (R1), we used a bootstrap procedure (Efron and Tibshirani 1994). We repeated the logistic-regression analysis 1,000 times by resampling subjects with replacement. This allowed us to estimate the confidence intervals for each cure given the data. Results showed that only the properties of the visual white matter projecting to fovea (R1 Figure 4A, blue) significantly predicted subjects’ class membership – the 95% CI were above chance level (50%) in the core white matter for R1 (Figure 4A blue horizontal bar).

**Figure 4.**
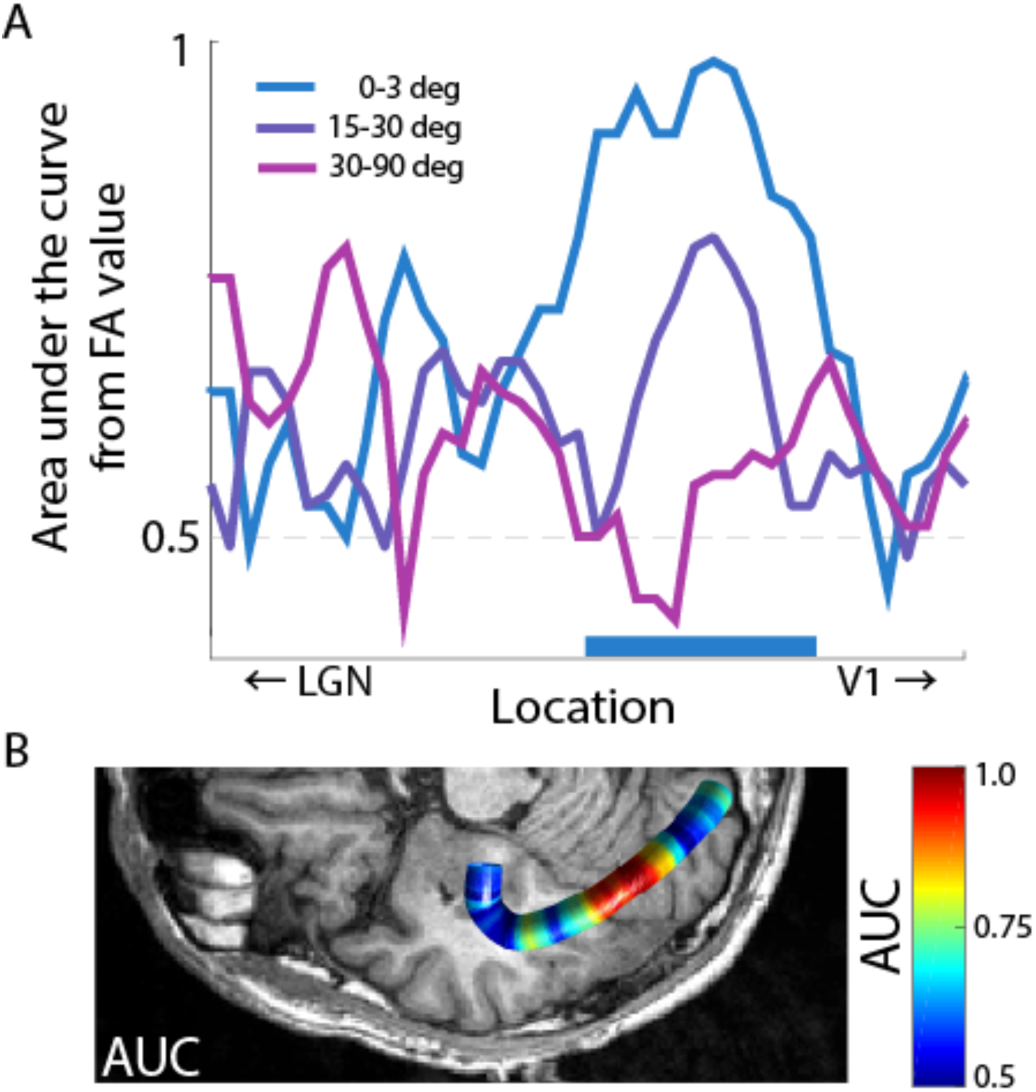
White matter along the optic radiation predicts subjects class membership. **A**. *Horizontal axis.* Spatial position along the optic radiation. *Vertical axis.* Area under the curve (AUC) of a logistic analysis in which FA values from different locations in the optic radiation were used to classify subjects as patients and controls. A value of 0.5 indicates no classification. A value of 1 indicates 100% classification performance; each subject is correctly classified in the correct category using FA values at that location of the OR. **B**. Anatomical localization of the measurements in **A**. Anatomical localization of the AUC values. AUC is plotted as function of location along the OR.

## Discussion

We presented an advanced anatomical characterization of the relation between white matter tissue-microstructure in the optic radiation and optic tract of patients affected by age-related macular degeneration. Our analyses, to our knowledge, are the first to demonstrate a conserved anatomical relation between the location of retinal damage (scotoma) and retinotopic-specific damage to white matter fascicles within the OR (Dougherty et al. 2005; Levin et al. 2010). Additionally, we report that the extent of visual impairment (behavioral measurement of visual acuity) in AMD patients is associated to the magnitude of white matter damage within the optic radiation. Hernowo and colleagues (2014) recently reported a reduction in the volume of the optic radiation using voxel-based morphometry in both juvenile and age-related macular degeneration (Hernowo et al. 2014b; Prins et al. 2016c). The present results build upon and extend these previous reports. In addition, our results are also consistent with a series of recent report demonstrating that the basic anatomy, network organization of the visual field maps in V1 is unaffected in individuals with macular degeneration (H. Baseler et al. 2010; H. A. Baseler et al. 2011; Shao et al. 2013; Haak et al. 2016a). We demonstrate that the effect of macular degeneration has very specific and anatomically localized effects on the visual white matter and that the magnitude of these effects predict visual acuity. Importantly, we show that such effects are robust and can be measured using in vivo method and magnetic resonance imaging data with clinical quality when such data is associated with advanced anatomical characterization.

We identified both the optic tracts and chiasms in every subject and found no significant decrease in FA in either of these structures. There are at least two reasons for such small changes in pregeniculate white matter: (1) there is no effect or (2) our measurements do not provide sufficient resolution to approach these structures. The human optic tract is a small structure on the order of 2-3 *mm* in diameter. It is embedded in an area with complex morphology where corticospinal fluid and supportive tissue types are intermixed (Parravano, Toledo, and Kucharczyk 1993). Our advanced dMRI measurements map brain tissue at 2 *mm* resolution. For this reason, it is likely that signals from the optic tract mixed with those of nearby tissues hiding any possible effect of AMD to the optic tract. We speculate that given the retinal damage due to AMD and our reports of associated damage to post-geniculate white matter damage to pregeniculate white matter is highly likely. Future higher-resolution studies might be able to report such damage. Our results are consistent with recent reports demonstrating that the white matter within the optic radiation is affected by AMD (Malania et al. 2017). We go beyond these previous reports by utilizing an anatomically-informed method that allowed us to characterize the specificity of the relation between locations of retinal damage and the visual white matter.

Our results extend a series of reports demonstrating a pervasive effect of eye disease on the visual white matter (Ariel Rokem et al. 2016; Millington, Ajina, and Bridge 2014). The biological mechanisms relating eye disease and the visual white matter are not completely understood. Indeed, the white matter could be affected by eye disease in at least several ways. For example, retrograde and anterograde degeneration (such as Wallerian degeneration (Pierpaoli et al. 2001; Rotshenker 2007)) could all be factors affecting the visual white matter depending the type of disease. The current understanding of the etiology of macular degeneration, is that it might be due to abnormal neo-vascularization that causes the death of photoreceptors and it is likely confined within the retinal tissue (Mandai et al. 2017a). This suggests a possible mechanisms for transsynaptic effects from the photoreceptors to the brain tissue. Indeed, our current results, previous ones (Malania et al. 2017) as well as results demonstrating a reduction in cortical volume in AMD (Hernowo et al. 2014a; Prins et al. 2016) all provide strong convergent evidence that cell death within the retina could affect the brain tissue via transsynaptic degeneration (Prins, Hanekamp, and Cornelissen 2016; Haak et al. 2016b).

The importance of precision approaches to the study of the human brain has been recently highlighted (Dubois and Adolphs 2016). We demonstrate the importance of informed anatomical characterization, methodology and study design in clarifying primary mechanism in the human white matter (Wandell 2016; Devlin and Poldrack 2007). We also show that beside easy to give away criticisms (Thomas et al. 2014), dMRI and fiber tractography offer much to inform scientists about fundamental mechanisms of the brain. In our case, the careful combination of *post mortem* anatomy, with cutting edge *in vivo* computational neuroanatomy methods provided convergent knowledge to design experiments helpful to characterize very specific effects of neurodegeneration in the human brain. The magnitude of the effect reported here in predicting brain and behavior effects from eye disease is enhanced by the fact that the results can be measured even at the clinical resolution of the data we used. There is much to learn about the human brain and convergent evidence from diverse measurements, models and statistical methods is paramount for discovery (Pestilli 2015; Ling, Jehee, and Pestilli 2015; Pestilli 2018).

## The visual white matter for predicting behavioral function, patient diagnosis and clinical intervention

A large proportion of AMD cases are believed to be neurovascular in nature (Wood et al. 2000; Jager, Mieler, and Miller 2008; Fine et al. 2000). For this reason, anti-vascular endothelial growth factor (VEGF) therapy is the most widely used form of treatment for AMD. Recently, modern intravitreal anti-VEGF therapy has improved treatment of AMD. Yet, unfortunately, not all participants respond to these therapies. A mere 40% of patients regain full visual function after intravitreal anti-VEGF therapy (D. M. Brown et al. 2006; Gonzalez and C 2011; Heier et al. 2012). To date, it is not clear why different AMD patients respond differently to this treatment. Measurements of white matter damage as a result of AMD could, in principle, be used to differentiate patients in different subgroups. The capacity to characterize different subtypes of patients given the extent of white matter damage could, in turn, improve the prediction power of therapeutic efficacy of treatments, such as intravitreal anti-VEGF therapy and novel iPS (induced pluripotent stem) cells transplant therapy (Mandai et al. 2017b), for refractory retinal diseases.

A number of studies have recently reported changes in white matter microstructure along the primary visual pathways as a result of ophthalmic eye disease (see (Prins, Hanekamp, and Cornelissen 2016c; H. D. H. Brown et al. 2016; Wandell 2016; Ariel Rokem et al. 2017) for reviews). Major examples, of impaired white matter tissue microstructure within the primary visual pathways have shown that retinitis pigmentosa (Pan et al. 2007), Leber hereditary optic neuropathy (Barcella et al. 2010; Milesi et al. 2012; Rizzo et al. 2012; Ogawa et al. 2014), amblyopia (Allen et al. 2015b; Duan et al. 2015b), glaucoma (Dai et al. 2013; Engelhorn et al. 2012; Garaci et al. 2009; Lee et al. 2014; Hernowo et al. 2011) and optic neuritis (Ciccarelli et al. 2005; Sergott 2011; Li et al. 2009) can affect white matter microstructure estimated *in vivo* using either diffusion-weighted magnetic resonance imaging methods or volumetric analyses. The current work adds to the growing evidence of strong ties between a variety of eye disease and corresponding major effects on brain tissue. It suggests that eye diseases might cause a generalized change in brain tissue properties. Future studies will be necessary to identify the extent to which damage to brain tissue microstructure due to eye disease goes beyond effects in the early visual pathways. Clarifying the extent of damage to white matter beyond the early visual pathways will have the potential to help identify differences between groups of patients and clarify reasons for differences in the effectiveness of treatments.

## Supplementary information

### Fractional anisotropy decreases in the optic tract and radiation

**Supplementary Figure 1.**
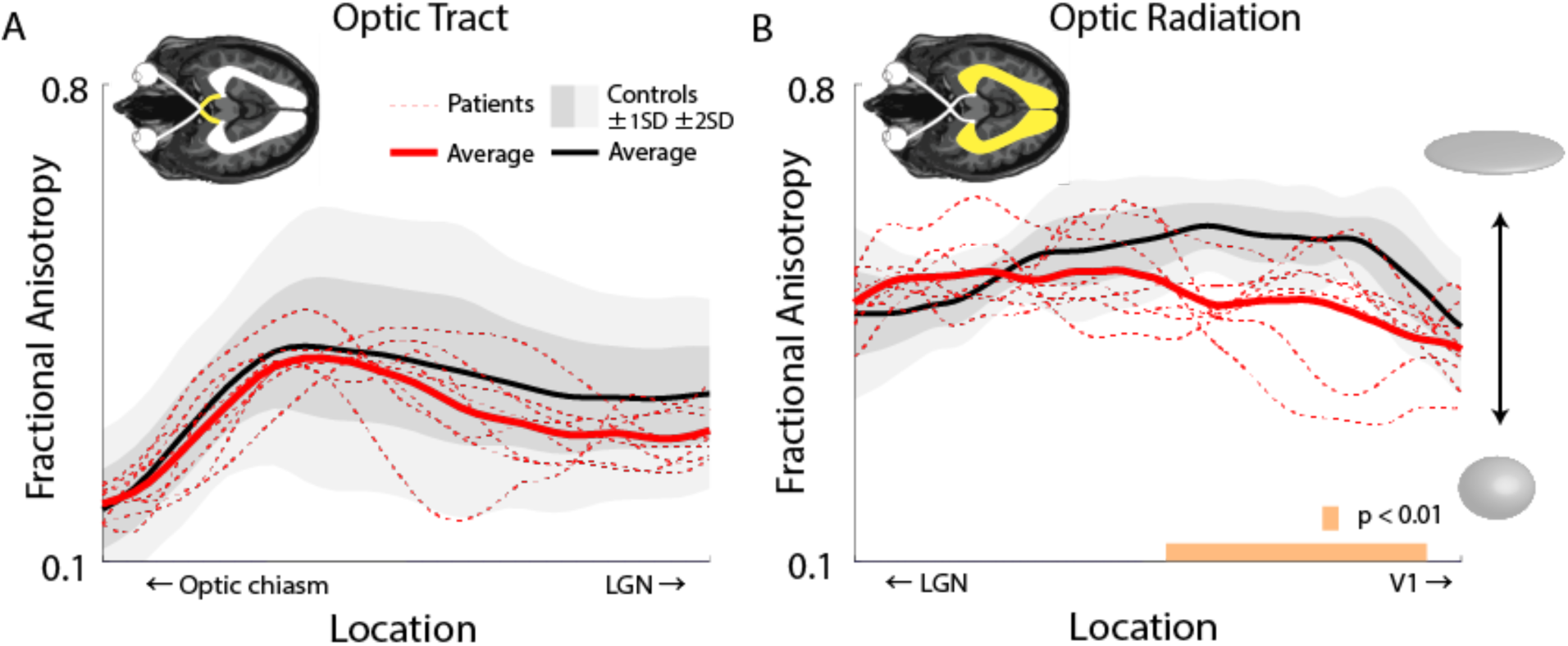
White matter tissue microstructure, fractional anisotropy, along the optic tract and radiation. **A**. Measurements of FA as function of anatomical position along the optic tract (OT). The x-axis represents the anatomical position along the optic tract, going from the optic chiasm (left) to the lateral geniculate nucleus (LGN, right). The y-axis shows fractional anisotropy, FA, measurements. In black is the average FA across control subjects, with dark and light gray shading indicating ± 1 and 2 standard deviations, respectively. In red measurements of FA in AMD patients, with individual measurements (dashed) and group mean (continuous). Measurements were averaged between hemispheres. **B**. Measurements of FA as function of anatomical position along the optic radiation (OR). Plot conventions similar to **A**, where the leftmost portion of the graph is closest to the LGN and the rightmost portion to primary visual cortex (V1). Orange bar shows locations with statistically significative difference between groups (unpaired t-test p < 0.01). Ellipsoid describes visually the relation between FA and the shape of the diffusion tensor model (P. J. Basser and Pierpaoli 1998; Peter J. Basser and Jones 2002).

**Supplementary Figure 2.**
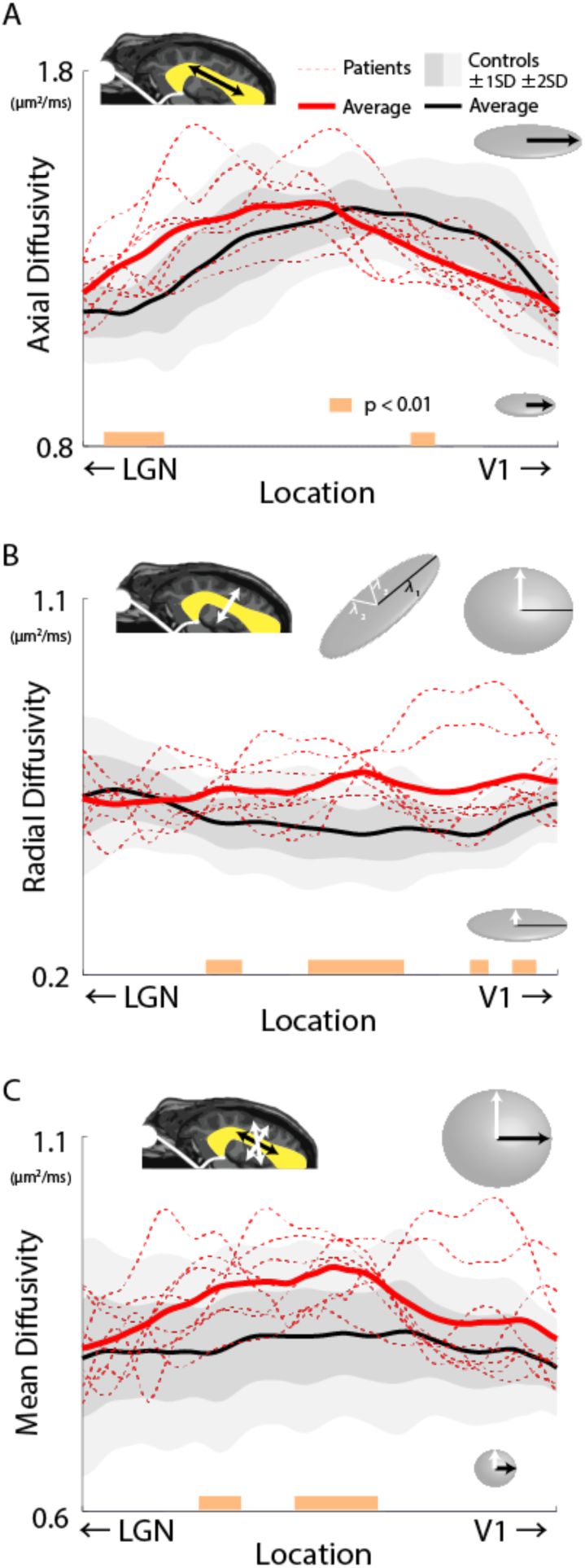
A, B and C. Additional white matter tissue microstructure estimates along the optic radiation. **A**. Tract profile of the optic radiation (OR) showing axial diffusivity for patients and the control group. **B**. Tract profile of the optic radiation showing radial diffusivity for patients and the control group. **C**. Mean diffusivity along the optic radiation. Differences in MD are evident across the central portion of the OR. Insets show the approximate relation between measurements in **A**, **B** and **C** and OR anatomy, as well as the nature of the measurements in relation to the tensor model (P. J. Basser & Pierpaoli, 1998).

**Supplementary Figure 2D.**
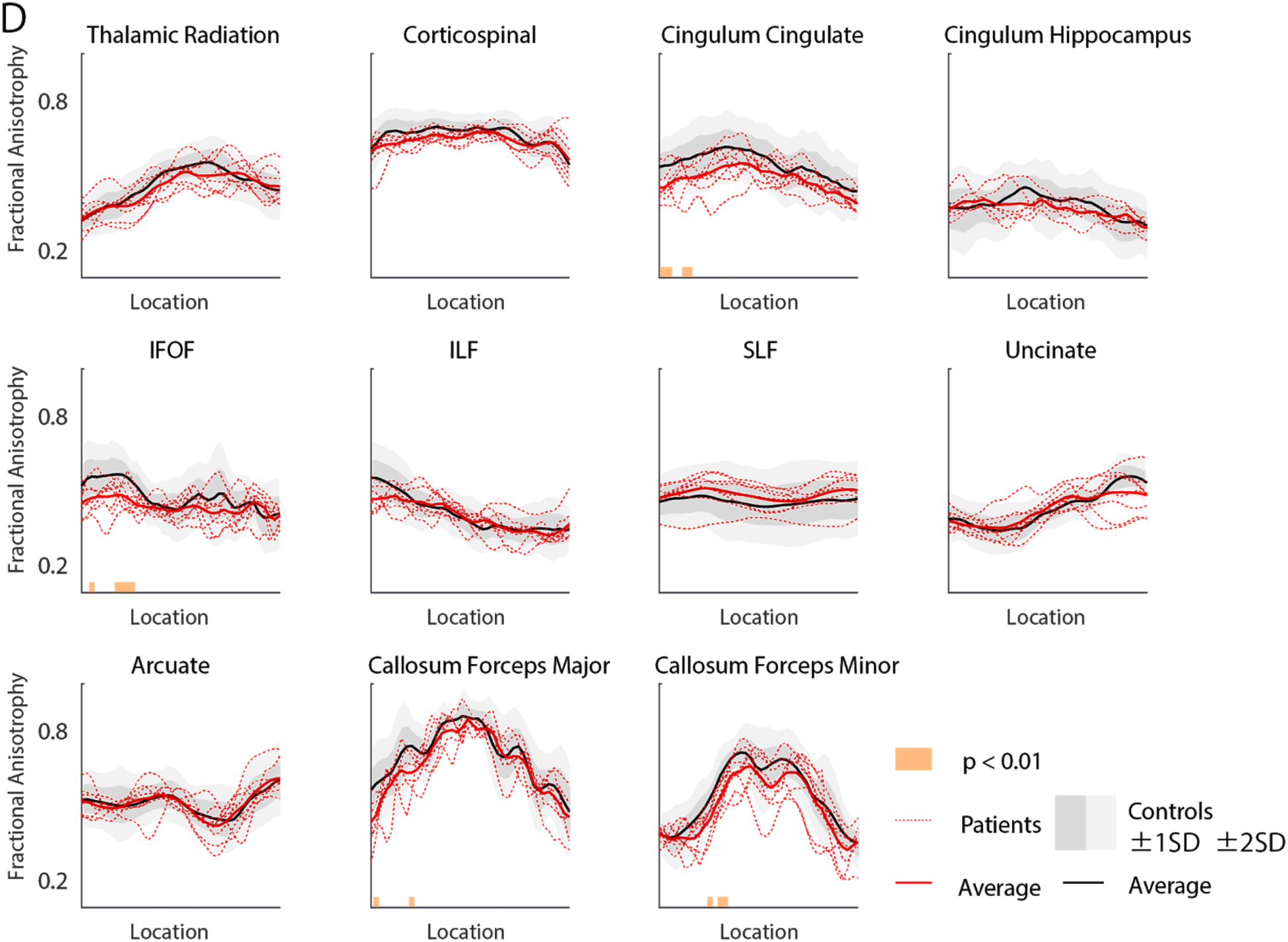
Analysis of the major human white matter tracts. To test whether the differences in FA found in the central portion of the OR only are not due to general differences in the brain white matter between patients and controls, we analyzed eleven major human white matter tracts (Mori et al. 2005; Yeatman et al. 2012). We found that, out of 440 tested white matter locations (nodes in the tract profiles), only 15 locations had a significant difference in FA between patients and controls (p<0.01). This is a small number of locations given the large number of comparisons performed. The results of this analysis is interpreted to demonstrate that although some small statistical differences between the two groups can be found, this is a small difference, is likely due to statistical factors (expected significance by chance given the large number of comparisons). This analysis demonstrate that the difference in the FA of the OR cannot be due to generalized difference the white matter between controls and patients.

**Author Contributions**
Designed the study: SO HT TN YM. Performed the experiments: SY SO HH MT AM YM. Analyzed the data: SY SO HH FP. Institutional support: KM. Wrote the paper: SY SO HH FP.

## Acknowledgements

We thank Brian A. Wandell, Sophia Vinci-Booher, Brent McPherson and Bradley Caron for comments on an early version of the manuscript. Takaaki Hayashi for clinical evaluation of the patients. Ikuya Murakami for institutional support. YS is supported by JSPS Grant-in-Aid for Young Scientists (B) 80570332. SO is supported by JSPS Grant-in-Aid for Young Scientists (B) 17K18131. HH is supported by JSPS Grant-in-Aid for Young Scientists (B) 26870605. YM is supported by The Jikei University Research Fund. F.P. is supported by NSF IIS-1636893, NSF BCS-1734853, NIH NIMH ULTTR001108, a Microsoft Research Award and the Indiana University Areas of Emergent Research initiative “Learning: Brains, Machines, Children,” and Pervasive Technology Institute.

## Notes

**Disclosure of potential conflicts of interest.** Authors declare no conflict.

